# Blueprint for Phasing and Assembling the Genomes of Heterozygous Polyploids: Application to the Octoploid Genome of Strawberry

**DOI:** 10.1101/2021.11.03.467115

**Authors:** Michael A. Hardigan, Mitchell J. Feldmann, Dominique D.A. Pincot, Randi A. Famula, Michaela V. Vachev, Mary A. Madera, Philipp Zerbe, Kristin Mars, Paul Peluso, David Rank, Shujun Ou, Christopher A. Saski, Charlotte B. Acharya, Glenn S. Cole, Alan E. Yocca, Adrian E. Platts, Patrick P. Edger, Steven J. Knapp

## Abstract

The challenge of allelic diversity for assembling haplotypes is exemplified in polyploid genomes containing homoeologous chromosomes of identical ancestry, and significant homologous variation within their ancestral subgenomes. Cultivated strawberry (*Fragaria × ananassa*) and its wild progenitors are outbred octoploids (2n = 8x = 56) in which up to eight homologous and homoeologous alleles are preserved. This introduces significant risk of haplotype collapse, switching, and chimeric fusions during assembly. Using third generation HiFi sequences from PacBio, we assembled the genome of the day-neutral octoploid *F. × ananassa* hybrid ‘Royal Royce’ from the University of California. Our goal was to produce subgenome-and haplotype-resolved assemblies of all 56 chromosomes, accurately reconstructing the parental haploid chromosome complements. Previous work has demonstrated that partitioning sequences by parental phase supports direct assembly of haplotypes in heterozygous diploid species. We leveraged the accuracy of HiFi sequence data with pedigree-informed sequencing to partition long read sequences by phase, and reduce the downstream risk of subgenomic chimeras during assembly. We were able to utilize an octoploid strawberry recombination breakpoint map containing 3.6 M variants to identify and break chimeric junctions, and perform scaffolding of the phase-1 and phase-2 octoploid assemblies. The N50 contiguity of the phase-1 and phase-2 assemblies prior to scaffolding and gap-filling was 11 Mb. The final haploid assembly represented seven of 28 chromosomes in a single contiguous sequence, and averaged fewer than three gaps per pseudomolecule. Additionally, we re-annotated the octoploid genome to produce a custom *F. × ananassa* repeat library and improved set of gene models based on IsoSeq transcript data and an expansive RNA-seq expression atlas. Here we present ‘FaRR1’, a gold-standard reference genome of *F. × ananassa* cultivar ‘Royal Royce’ to assist future genomic research and molecular breeding of allo-octoploid strawberry.

## Introduction

Until recently, breeding and genetics research of garden strawberry (*Fragaria × ananassa* Duchesne ex Rozier) and its wild relatives was challenged by their complex allooctoploid genomes (2n = 8x = 56). The publication of the first chromosome-scale genome assembly of an octoploid strawberry (1) rapidly enabled studies that have dramatically increased our understanding of octoploid subgenome evolution (1, 2), subgenome diversity and domestication (3, 4), and the genetic basis of traits such as resistance to economically destructive pathogens (5). Notably, these studies have shown that many diploid approaches to breeding, quantitative genetic, and population genetic research can be directly applied to the four diploid-behaving subgenomes of *F. × ananassa*, so long as haploid assemblies of the octoploid physical genome with fully separated homoeologs are present.

While the Camarosa v1 genome assembly dramatically improved our ability to conduct strawberry genetic studies, there remained room for improvement in the effort to develop a reference-quality genome sequence. The Camarosa v1 genome shows the phase-switching, limited contiguity, and local assembly errors expected from a short-read assembly of a heterozygous polyploid. Other features we wished to address included a number of megabase (Mb)-scale scaffolding errors (inversions and translocations), retention of homologous allelic contigs within homoeologous haploid pseudo-molecules, non-uniform orientations of homoeologous pseudomolecules, and a chromosome nomenclature that does not reflect the proposed origins of the respective subgenomes. For this strawberry genome assembly project, we used the heterozygous day-neutral cultivar ‘Royal Royce’ released in 2020 by the University of California, Davis strawberry breeding program. We re-approached the problem of octoploid genome assembly with ‘Royal Royce’ using previously untapped resources: highly accurate long read (HiFi) data (6), pedigree-informed parental sequences to enable trio-binning, a recombination breakpoint map containing 3.6 M markers for identifying chimeras and accurate scaffolding (2), and an improved transcriptomic dataset for gene annotation that included IsoSeq long reads from ‘Royal Royce’ and an expanded panel of RNA-seq data from eighteen expression studies.

PacBio HiFi reads are generated using circular consensus calling over the same sequence, improving their base-calling accuracy to over 99% (6). This was important for confident assembly of an octoploid genome due to the risk of introducing subgenome collapses, fusions, or chimeras into the dataset during the all-vs-all alignment and error correction steps required for traditional long reads, where inter-subgenome alignments and homologous allelic sequence alignments can easily be conflated and used improperly as evidence for read “correction”. The accuracy of HiFi reads also means they can directly support trio-binning using Illumina sequences derived from parental genotypes. We used this approach in the current study, partitioning ‘Royal Royce’ HiFi sequences into two parent-specific haplotype bins, and subsequently conducting parallel assemblies with the respective haploid datasets. The length and accuracy of the HiFi data allowed us to enforce highly stringent thresholds for read alignment during assembly, in an effort to minimize the occurrence of subgenome chimeras. We were also able to use a recombination breakpoint map containing 3.6 M genetically mapped markers to identify subgenome chimeras in the resulting phased assemblies. While the use of stringent thresholds for HiFi read alignment during assembly did not fully prevent subgenome chimeras, they were sufficiently limited that we were able to identify chimeric junctions using the recombination breakpoint map and introduce breaks at these locations.

Both of the ‘Royal Royce’ haploid assemblies (phase-1 and phase-2) were approximately 784 Mb in size, with N50 values of 11 Mb before scaffolding and gap-filling, and N50 values of 23 Mb after scaffolding and gap-filling. Maximum contig sizes were 35 Mb, and included multiple single-contig chromosome assemblies. We performed *de novo* repeat annotation and gene annotation of both haploid assemblies, identifying nearly as many gene models as the 108k in Camarosa v1, at roughly 102k per haploid genome. The quality of gene models improved due to the use of IsoSeq full length transcript sequences for annotation. To support octoploid strawberry breeding and genetics research, we selected the best quality phase-1 or phase-2 pseudomolecule and corresponding gene models for each of the 28 haploid *F. × ananassa* chromosomes to construct a synthetic haploid genome of the cultivar Royal Royce, now called ‘FaRR1’, that significantly improves on the existing Camarosa v1 haploid assembly (1). In addition to minimizing haplotype-switching resulting from the assembly of heterozygous sequences, 15 of 28 chromosomes are represented by pseudomolecules containing two or fewer contiguous sequences. We have also updated the chromosome nomenclature in the FaRR1 assembly to represent the proposed diploid subgenome origins of each pseudomolecule as in wheat (e.g., A, B, C, D genomes). This assembly offers an improved tool for marker development, gene discovery, and functional genetics in a species where the identification and separation of homoeologous variation remains a critical challenge for targeting subgenome-specific causal loci.

## Results and Discussion

### Octoploid Strawberry Genomic and Transcriptomic Datasets

We obtained the published PacBio HiFi genomic sequence reads for *F. × ananassa* cultivar ‘Royal Royce’, generated as described by (6). The data was generated on the Sequel II platform and contained a total of 83 Gb of sequence in 3.8 M HiFi reads with an average size of 21.7 kb. We supplemented the ‘Royal Royce’ HiFi sequence data with Illumina paired-end 150 bp whole genome shotgun (WGS) sequences for the ‘Royal Royce’ parent genotypes ‘04C009P005’ and ‘05C099P006’ to support trio-binning. Both parent genotypes were sequenced to a depth of 120X coverage. We generated IsoSeq data for 15 ‘Royal Royce’ libraries representing 15 tissues and treatments (Supplementary Table 1). We supplemented the ‘Royal Royce’ IsoSeq data with publicly available RNA-seq datasets from 67 octoploid strawberry RNA-seq libraries derived from 18 previous studies and representing various tissues and treatments (Supplementary Table 2). In total, we obtained 12.1 Gb of IsoSeq sequence data and 508.0 Gb of short read RNA-seq data as input for transcriptome assembly and downstream annotation.

### HiFi Trio-Binning

We circumvented the bioinformatic hurdle of “unzipping” allelic contigs in an octoploid genome assembly containing a complex mixture of homologous and homoeologous alleles by trio-binning the ‘Royal Royce’ HiFi reads into parent-specific read pools, then performing parallel assembly of the parental haplotypes. PacBio HiFi reads should support accurate Illumina-based trio-binning due to their low error rates (base-call accuracy *≥* 99.5%) (6). We expected any resulting trio-binned read pools to contain homozygous parental sequence information in the sense that they represent distinct haploid chromosome sets (n=28), while still retaining allo-polyploid ‘fixed heterozygosity’ between the four ancestrally related chromosome sets within each haploid read pool. Trio-binning was supported by Illumina sequence datasets generated for the maternal parent (‘04C009P005’) and paternal parent (‘05C099P006’) genotypes. Hereafter, sequences and assemblies derived from the maternal parent bin are referred to as ‘phase-1’, and those derived from the paternal parent bin are referred to as ‘phase-2’. We used the methods and software published by (7) (https://github.com/esrice/trio_binning) to identify unique 21-nt k-mers in the parental short read sequence datasets, and for estimating the histogram of parental k-mer counts. Subsequently, we retained k-mers with counts ranging from 20-250 observations. The distribution of parent-specific k-mer frequencies in reads with at least 100 unique k-mers clearly highlighted the set of ‘Royal Royce’ HiFi reads representing the heterozygous regions between corresponding homologous chromosomes of the two parents (Fig 1). For partitioning of HiFi data into parental bins (trio-binning) we required HiFi reads to contain at least 100 parent-specific k-mers, and required that at least 80% of total unique k-mers were specific to the dominant parent. Reads that failed to meet these criteria were assumed to represent regions in which the parents shared identical-by-state (IBS) haplotypes. In total, 31% of HiFi reads were classified as representing parent-specific haplotypes from heterozygous genome regions. After trio-binning, the phase-1 read pool contained 70.8 Gb of sequence and the phase-2 read pool contained 69.8 Gb of sequence, providing approximately 80X haploid genome coverage in each dataset. Both read pools were filtered to retain approximately 40X coverage of reads greater than 22.5 kb. The average HiFi read length after filtering was 25.2 kb.

**Fig. 1.**
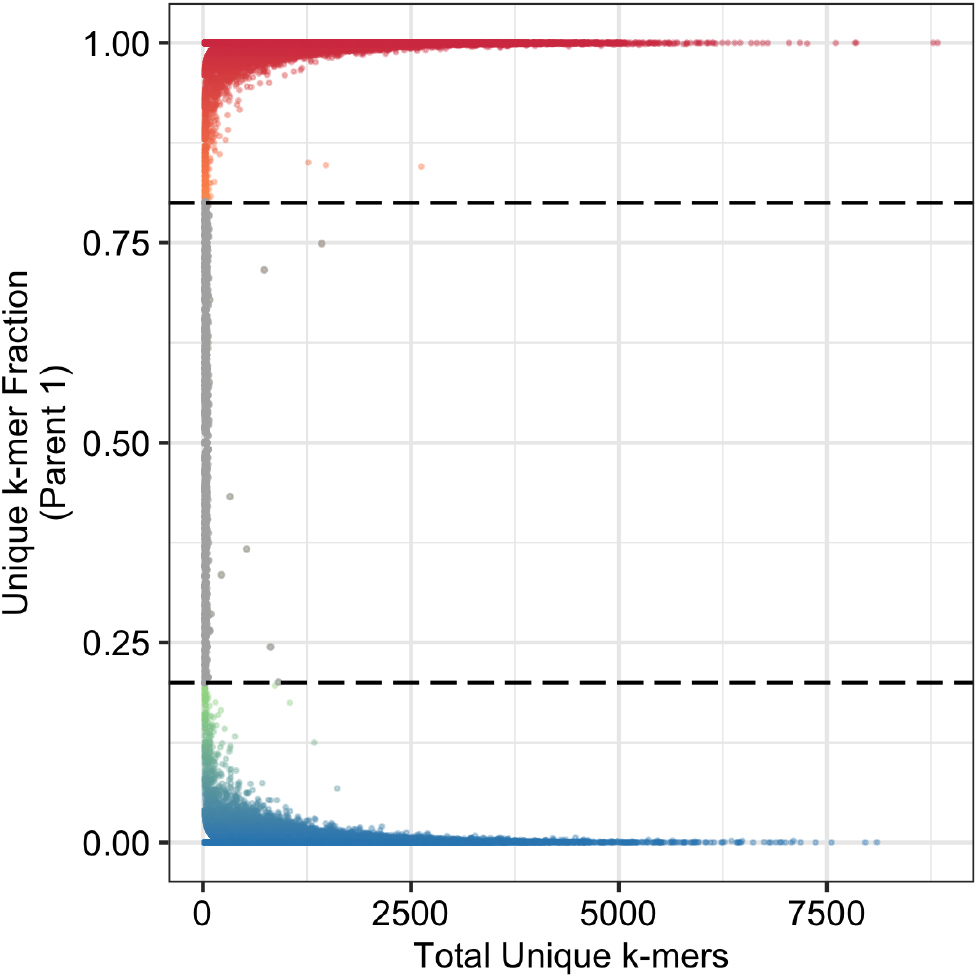
Distribution of total unique k-mers plotted against parent-specific k-mer frequency in the HiFi sequences used for ‘Royal Royce’ genome assembly.

### Phased Octoploid Genome Assembly

The phase-1 and phase-2 HiFi read pools were assembled in parallel using the software FALCON (https://github.com/PacificBiosciences/FALCON). Because of the high base-call accuracy of HiFi reads, we skipped the initial errorcorrection step used by FALCON to process long reads containing 5-15% error rates. Instead, the HiFi reads were designated as ‘pread’ input for FALCON and used directly to calculate sequence overlaps for contig assembly. Due to strawberry’s four ancestrally related subgenomes that increase the potential for chimeric assemblies, we required a minimum overlap length of 5,000 bp and minimum overlap identity of 99.5% for pread overlap detection. To further reduce the potential for mixing homoeologous or allelic preads, in the final assembly stage we used a 99.8% identity threshold for overlap filtering and instructed FALCON to retain the best subset of overlaps up to a cutoff of 15X depth for assembly, well below the 20-30X haploid sequence coverage available within any given subgenome.

The initial FALCON output contained 979 Mb across 310 contigs in the phase-1 assembly and 948 Mb across 272 contigs in the phase-2 assembly, roughly 150-200 Mb over the approximate haploid genome size of 800 Mb. We performed all-vs-all alignments of FALCON contigs within the phase-1 and phase-2 pools and found that in each pool greater than half of the assembled contigs were fully contained (100% query coverage) within a larger contig at a sequence identity above 99%, but below the 99.8% identity threshold that we enforced for pread overlap filtering. The mean basecall accuracy of HiFi reads ranged from 99.18-99.73% in the Hon et al. (2020) study which produced the ‘Royal Royce’ HiFi dataset (6). We concluded that setting a 99.8% identity threshold for pread overlap filtering, which straddles the approximate base-calling accuracy of the HiFi sequences, reduced the probability of subgenome chimeras while also yielding artifactual ‘allelic’ contigs representing regions in which error rates approached or exceeded 0.2%. We addressed this by using HaploMerger2 (8) on a chromosome-specific basis (56 independent contig pools) to collapse any contigs fully contained (100% query coverage, *≥* 99% identity) within a larger contig from the same chromosome. Below we describe our approach for using recombination break-point maps for chimera detection and then for binning the FALCON contigs into 56 chromosome-specific pools to collapse these artifacts.

We used a high-density recombination breakpoint map containing 3.6 M genetically mapped variants published by (2) to paint the assembled FALCON contigs by homoeologous chromosome origin, detect and split subgenome-chimeras in the FALCON contigs, then bin the FALCON contigs by chromosome origin for collapsing redundant sequence artifacts. For each of the 3.6 M genetically mapped variant sites containing octoploid linkage group information, we extracted the 2 kb sequence flanking its physical position in the Camarosa v1 assembly used for variant calling (2). We aligned these 2 kb anchoring sequences to the phase-1 and phase-2 FALCON contigs using BLAST, and used the best-scoring alignments to determine the corresponding physical positions of the genetically mapped markers on the ‘Royal Royce’ phase-1 and phase-2 contigs. We used the software ALLMAPS (9) to identify subgenome chimeras based on discordant genetic map positions or linkage group origins within the physical contigs. Despite the stringent alignment thresholds used to assemble HiFi reads, we identified five subgenome chimeras in the phase-1 contigs and four subgenome chimeras in the phase-2 contigs. Because of this relatively small number of subgenome chimeras, we manually inspected the chimeric phase-1 and phase-2 contigs to identify junctions where the anchored genetic markers transitioned between independent linkage group regions along the physical contig space. We manually broke the chimeric contigs at these junctions, using the outermost variant sites from the respective independent linkage group regions as borders for the newly split contigs. None of the chimeric contigs contained sequences corresponding to more than two independent linkage group regions, such that each split yielded exactly two new contigs from the initial chimeric contig.

After resolving subgenome chimeras in the initial FALCON contigs, we realigned the 3.6 M genetically mapped markers to the new set of contigs and partitioned the respective phase-1 and phase-2 contig pools into 56 chromosome-specific contig bins based on the dominant corresponding linkage group. For phase-1 contigs we successfully placed 99.1% of the total assembled sequence into 28 chromosome-specific bins. For phase-2 contigs we successfully placed 99.2% of the total assembled sequence into 28 chromosome-specific bins. We used LastZ and HaploMerger2 within each chromosome-specific contig bin to collapse any redundant sequence artifacts, which we defined as contigs fully contained within a larger contig (100% query coverage) at over 99% sequence identity (8). After breaking the chimeric assemblies and collapsing redundant sequence artifacts, the phase-1 assembly contained 136 contigs with a total size of 784.7 Mb and N50 of 11.0 Mb, while the phase-2 assembly contained 131 contigs with a total size of 784.0 Mb and N50 of 10.7 Mb (Table 1). These results indicate that enforcing strict alignment thresholds for HiFi overlap detection and filtering may improve the accuracy of polyploid subgenome assemblies, however, we were unable to fully prevent subgenome chimeras, and a byproduct of using alignment thresholds approaching 100% appears to be redundantly assembled contigs in multiple regions, despite the exclusion of allelic reads by triobinning.

**Table 1.** Assembly quality metrics for ‘Royal Royce’ phase-1 and phase-2 contigs before and after scaffolding and gap-filling steps and for the synthetic haploid assembly FaRR1.

| Assembly Metrics | Pre-Scaffolding |  | Scaffolding and Gap-Filled |  |  |
| --- | --- | --- | --- | --- | --- |
|  | Phase-1 | Phase-2 | Phase-1 | Phase-2 | FaRR1 |
| Number of Contigs | 136 | 131 | 81 | 77 | 65 |
| Number of Contigs $\geq 50$ kb | 134 | 129 | 79 | 75 | 62 |
| Genome Length (bp) | 784,724,251 | 784,042,449 | 784,481,303 | 783,923,278 | 786,542,606 |
| Genome Length for Contigs $\geq 50$ kb (bp) | 784,626,701 | 783,961,397 | 784,385,248 | 783,842,146 | 786,435,331 |
| Largest Contig (bp) | 27,612,319 | 31,257,307 | 35,982,586 | 34,899,819 | 35,982,586 |
| GC Content (%) | 39.40 | 39.38 | 39.39 | 39.37 | 39.38 |
| N50 (bp) | 11,038,762 | 10,737,681 | 22,855,991 | 22,554,989 | 25,978,654 |
| N75 (bp) | 7,133,475 | 6,844,057 | 16,080,594 | 19,378,671 | 19,152,627 |
| L50 | 23 | 23 | 15 | 15 | 14 |
| L75 | 44 | 45 | 25 | 24 | 23 |

### Assembly Scaffolding and Polishing

We ordered and oriented the phase-1 and phase-2 contigs and performed genome scaffolding using the software ALLMAPS (9), employing the same recombination breakpoint map used previously for identifying chimeras and binning contigs by linkage group. As input, we provided the previously described information from 3.6 M genetically mapped variant sites and their physically anchored positions in the unscaffolded phase-1 and phase-2 contigs. In total, 99.5% of phase-1 contig sequence was scaffolded into 28 pseudomolecules and 99.5% of phase-2 contig sequence was scaffolded into 28 pseudomolecules. Thus, nearly all parental-derived chromosome sequence was represented in the 56 chromosome-scale pseudomolecules. After scaffolding we performed gap-filling using phase-specific HiFi reads in both haploid assemblies with the software LR_Gapcloser (10). With this approach we closed 17 of 82 gaps in the phase-1 assembly pseudo-molecules and closed 20 of 81 gaps in the phase-2 assembly pseudomolecules. We then performed consensus-based error-correction of the phase-1 and phase-2 assemblies using the software Pilon (11). As input, we provided alignments of phase-specific HiFi reads as well as the corresponding parent-specific Illumina sequence reads used for trio-binning. Because each parent was a heterozygous individual, we excluded any parental Illumina read alignments containing more than a single mismatch, to prevent polishing based on reads derived from parental alleles that were not inherited by ‘Royal Royce’. Using this approach, we corrected 4,578 SNPs and 11,019 indels in the phase-1 assembly, and corrected 4,549 SNPs and 10,973 indels in the phase-2 assembly. We expected this larger proportion of indel errors in the assemblies due to the indel-heavy error profiles of PacBio sequences. The polished pseudomolecules were oriented identically relative to their corresponding diploid strawberry (*Fragaria vesca*) pseudomolecule in the v4.0 genome assembly (12).

From each of the 28 parental pairs of octoploid strawberry chromosomes we selected the most contiguous pseudomolecule from the corresponding phase-1/phase-2 parents sets to produce an optimal haploid genome assembly labelled ‘FaRR1’ that is suitable for calling heterozygous subgenome-level DNA variants in octoploid strawberry. The FaRR1 assembly contains 786.5 Mb of sequence and 28 chromosome-scale pseudomolecules accounting for 99.5% of the assembled genome, and 21 unanchored contigs which could not be assigned to a particular chromosome. Seven of the FaRR1 pseudomolecules (25%) are represented by a single contiguous sequence without gaps, and the FaRR1 pseudomolecules contained an average of 2.5 contigs (post-gap filling), indicating a high degree of contiguity in the final haploid assembly (Table 1).

### Octoploid Chromosome Nomenclature

The first chromosome-scale genome assembly of octoploid strawberry, Camarosa v1, inherited its homoeologous chromosome identifiers from the (13) genetic map utilized for genome scaffolding in that study (1). These chromosome identifiers were not associated with any particular ancestral subgenome origin. As a result, homoeologous chromosome suffixes (e.g., Fvb1-1, -2, -3, -4) did not specify similar phylogenetic origins within any of the four sets of seven chromosomes sharing a common suffix in the Camarosa v1 assembly. Given the differences in subgenome dominance, gene expression, and historically asymmetric DNA transfer between homoeologs, these identifiers are important for breeding and genetic studies. (3) published an updated octoploid strawberry chromosome nomenclature which utilized phylogenetic species assignment of subgenome transcript assemblies to reassign chromosomes into an A subgenome derived from *F. vesca*, a B subgenome derived from *F. iinumae*, and C and D subgenomes which shared the greatest degree of similarity with extant diploid species *F. nipponica* and *F. viridis*, respectively (3). Chromosome names in the FaRR1 genome assembly were updated from the Camarosa v1 nomenclature so that homoeologous chromosome identifiers in the octoploid genome now reflect the proposed diploid origins of each respective subgenome (A, B, C, D) (Supplementary Table 3).

### Assembly Quality and Completeness

We estimated the completeness of the FaRR1 assembly by BUSCO analysis of the core eudicot gene set (14). For comparison, we performed parallel analyses of the diploid *F. vesca* v4.0 assembly (12), as well as the four respective seven-chromosome subsets of FaRR1 representing its diploid-derived A, B, C and D subgenomes (Fig 2). We observed 98.1% complete gene models in both the octoploid and diploid strawberry genomes, indicating a high degree of completeness in the FaRR1 assembly. A majority of the complete gene models were duplicated in *F. × ananassa*, indicating that in almost every case there were unfragmented core eudicot genes on two or more subgenomes. However, the fraction of complete versus fragmented or missing core genes on the respective *F. × ananassa* subgenomes followed their dynamics of subgenome dominance: A > B > C D. In addition to assessing FaRR1 genome quality based on assembly of the core eudicot gene space, the assembly quality was also assessed using the LTR Assembly Index (LAI) (15). The LAI metric reflects the overall assembly continuity based on analysis of intact and total LTR retrotransposons. The LAI scores for ‘Royal Royce’ were above 18 for each subgenome on the phase-1 and phase-2 haplotypes, which is higher than the LAI obtained for Camarosa v1 as well as diploid *F. vesca* Hawaii 4 (Supplementary Table 4) (12). The LAI scores ranged from 18.2 to 19.33. These scores would classify FaRR1 at the high end of Reference quality and near goldstandard, given that (15) showed an average LAI score of 15.5 in bacterial artificial chromosome (BAC) derived assemblies. This is particularly impressive given that the ‘Royal Royce’ assembly is completely haplotype-phased and highly polyploid. Most plant genome that are considered to be gold-standard are diploid and only have a single haplotype assembled. We visualized whole-genome alignments of FaRR1 against the diploid *F. vesca* v4.0 assembly to compare their genome structures (Fig 3). For each of the seven ancestral diploid chromosomes, the respective octoploid homoeologs maintained a high degree of collinearity, with the exception of a major translocation on 2C. We also observed a region with significant structural variation relative to *F. vesca* on the distal arm of the four octoploid chromosome 1 homoeologs, including subgenome A. Finally, we plotted physical positions in the FaRR1 and Camarosa v1 genome assemblies against genetic positions in a random subset of 300,000 variants from the recombination map used previously (Fig 4). In addition to introducing conserved chromosome orientations, we observed that FaRR1 was more collinear with the recombination map than Camarosa v1 and resolved major scaffolding errors on multiple chromosomes: 2C, 2D, 4A, 6C. Furthermore, we observed that FaRR1 addressed cases in which the Camarosa v1 assembly retained and scaffolded interleaved allelic contigs. This was visible on chromosomes 7A and 7C; comparison to the recombination map revealed Camarosa v1 harbored overlapping allelic assemblies spanning large segments of the pseudomolecules. Based on these analyses, we concluded the FaRR1 assembly has a high degree of completeness and has helped to address scaffolding errors and cases of retained allelic contigs from previous octoploid assemblies.

**Fig. 2.**
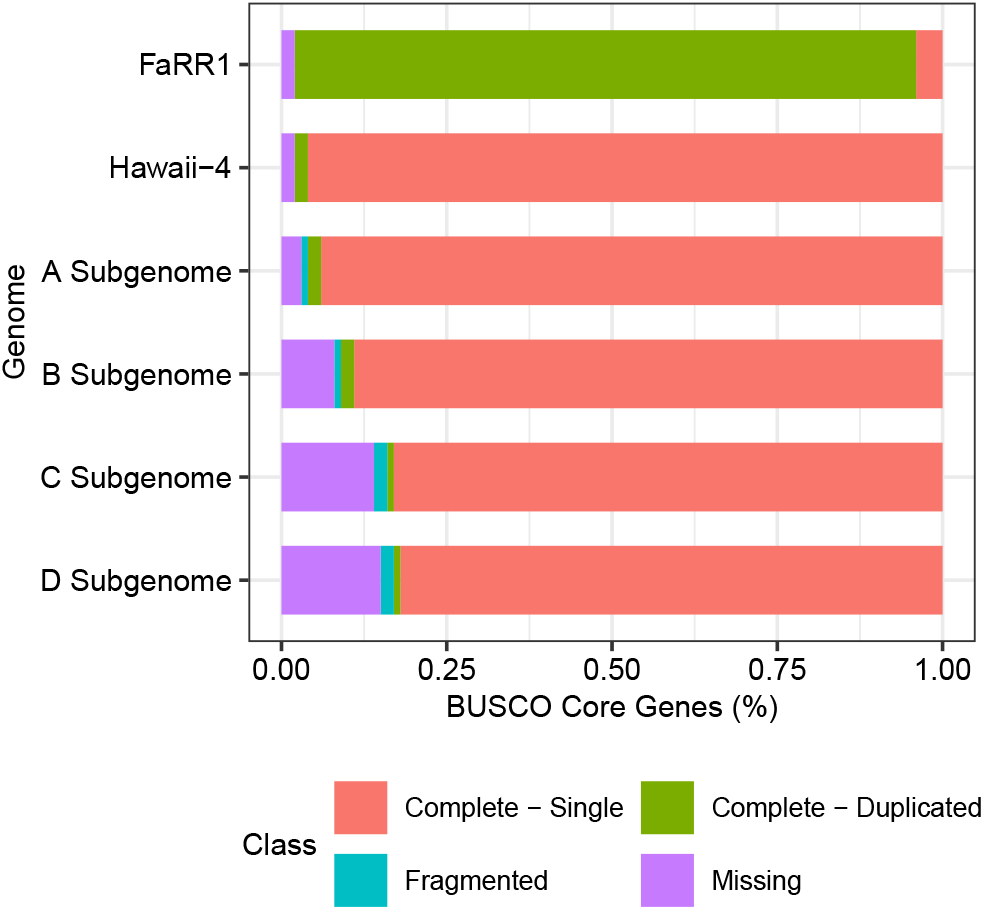
BUSCO analysis of FaRR1 genome assembly quality. Displays the relative fraction of core eudicot plant genes that are complete, fragmented, and missing in the octoploid *F. × ananassa* genome (FaRR1 v1.0), the diploid *F. vesca* genome (Hawaii-4 v4.0), and the four ancient diploid subgenomes contained within *F. × ananassa*.

**Fig. 3.**
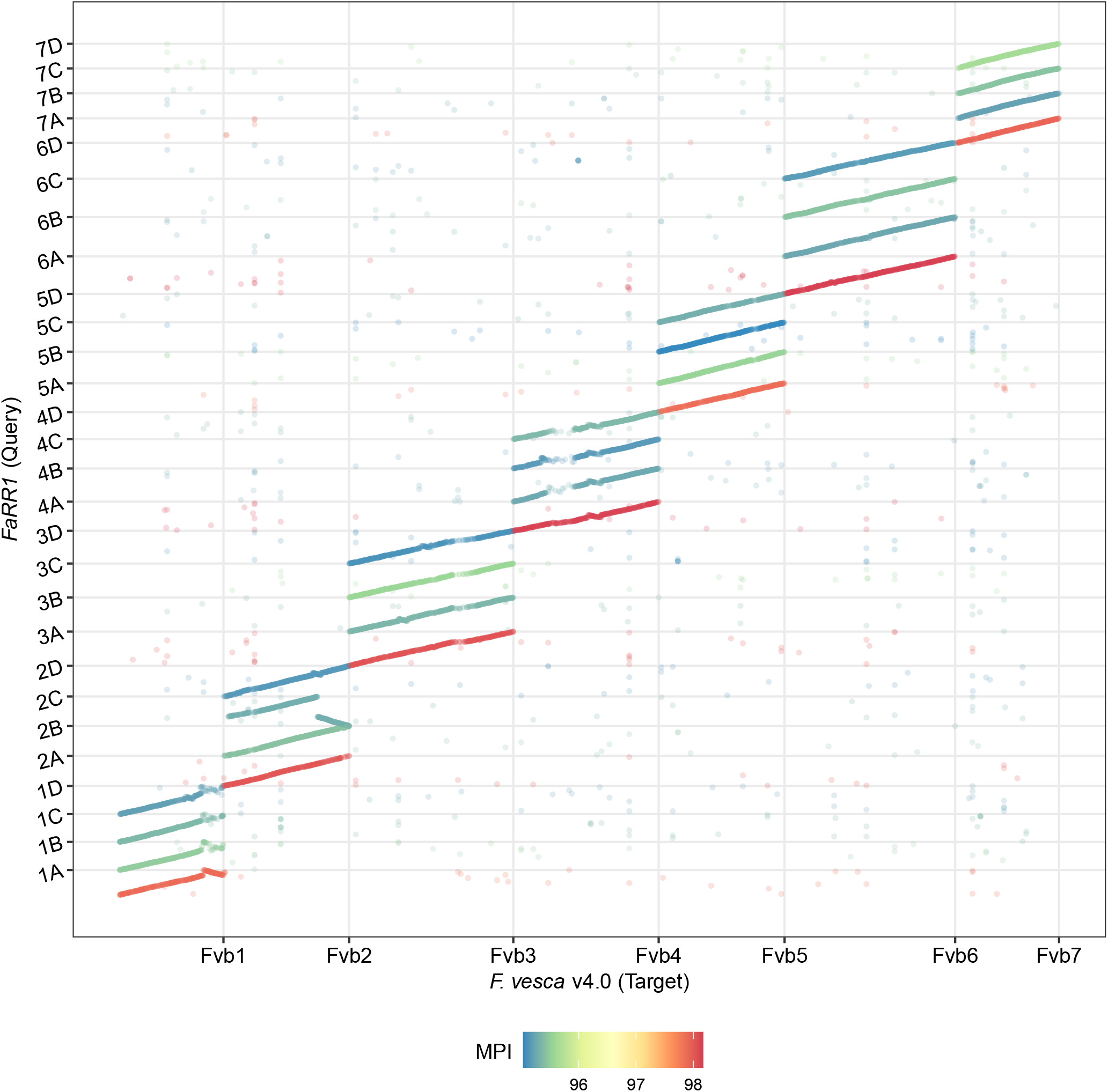
Dotplot visualization of whole-genome alignment of the octoploid FaRR1 v1.0 genome assembly to the diploid *F. vesca* genome assembly (v4.0). MPI is the mean percent identity of the query (FaRR1 v1.0) to the target (*F. vesca* Hawaii-4 v4.0).

**Fig. 4.**
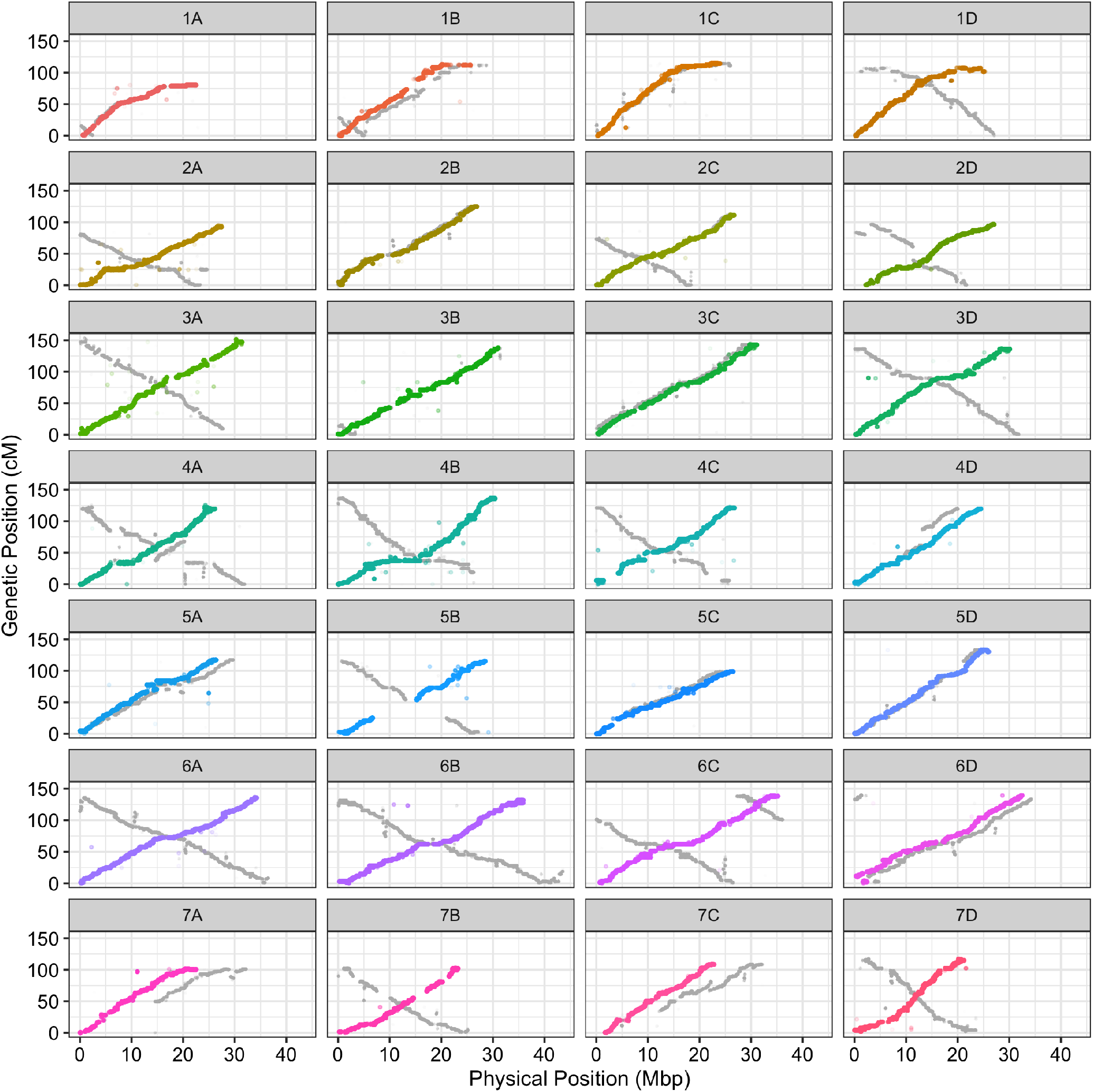
Genetic positions from a random subset of 1.7M markers from an octoploid strawberry recombination breakpoint map plotted against the corresponding physical positions in the FaRR1 v1.0 genome assembly (colored points) or Camarosa v1 reference genome assembly (gray points).

### Genome Annotation

To support an improved FaRR1 genome annotation, we generated an improved *F. × ananassa* transcriptome assembly containing the broadest possible set of transcripts from ‘Royal Royce’ and publicly available *F. × ananassa* expression data, with minimal redundancy. We produced IsoSeq data from 15 ‘Royal Royce’ tissue and treatment libraries (Supplementary Table 1), in addition to RNA-seq data from 67 broad *F. × ananassa* tissue and treatment libraries (Supplementary Table 2). The IsoSeq and RNA-seq datasets were aligned to the combined phase-1 and phase-2 genome assemblies. These alignments were used to perform reference-guided transcriptome assemblies for each library with Stringtie2 (16). The resulting reference-guided transcriptome assemblies were used as input for the program Mikado (17), which we used to parse all transcriptome assemblies by genome region, select the best transcript evidence at each locus, and attempt to split fused transcripts. As additional evidence to improve transcript scoring, we provided Mikado with a list of well-supported splice junctions detected from alignment data using Portcullis (18), protein homology information generated by BLAST alignment against the UniProt database (19), and open reading frame (ORF) scores generated using TransDecoder (20). We also provided Mikado with a custom scoring file to automatically prioritize transcript evidence derived from IsoSeq data in cases where both IsoSeq and RNA-seq assemblies were available. We obtained a non-redundant set of 160,375 transcripts after filtering to remove assemblies with no associated PFAM domains and ORFs shorter than 75 amino acid residues, yielding roughly 80,000 transcripts per haploid genome.

We generated a custom repeat library for *F. × ananassa* using the FaRR1 genome assembly and the ‘Extensive *de novo* TE Annotator’ (EDTA) pipeline, in order to identify transposable elements (TEs) and other repetitive sequences to avoid during gene annotation. In total, 38.42% of the combined phase-1 and phase-2 assemblies were annotated as repetitive sequence, with a majority of this repeat sequence contributed by LTR class transposable elements (Table 2). We performed gene annotation using the MAKER pipeline (21). This included an initial round of evidence alignment, followed by two rounds of training and prediction using of the SNAP and Augustus gene prediction programs. In the first round we provided the EDTA custom repeat library as masking evidence, the non-redundant set of Mikado transcript assemblies as EST evidence, and UniProt plant proteins (19) as protein evidence. Resulting gene models with proteins longer than 75 amino acid residues and AED scores below 0.20 were retained as evidence to train SNAP and Augustus to perform gene prediction for two additional rounds. MAKER predicted 248,061 total gene models, roughly 124,000 models per haploid genome. The total annotated gene space accounted for 41% of the *F. × ananassa* genome. Next, we identified and excluded TE-related gene models by three methods: 1) using Hidden Markov models to scan gene model protein sequences for TE-related PFAM domains, 2) BLAST alignment of gene model protein sequences against a database of transposases (http://weatherby.genetics.utah.edu/MAKER/wiki/index.php/Repeat_Library_Construction-Advanced), and 3) BLAST alignment of gene model transcript sequences against DNA sequences in the *F. × ananassa* repeat library generated by the EDTA pipeline. In order to exclude fragmented genes that were not TE-related, we excluded gene models that both encoded proteins with fewer than 65 amino acid residues and had annotation edit distance scores greater than 0.70. The majority of excluded gene models (85%) were TE-related. A total of 203,170 gene models remained in the final annotation after excluding fragmented and TE-related gene models, with 101,721 models in the phase-1 assembly, 101,449 models in the phase-2 assembly, and 101,793 in the FaRR1 assembly. Thus, each haploid genome contained approximately 94% of the total number of annotated genes in Camarosa v1 (1). Comparison of the cumulative AED score distribution for representative transcripts in the Camarosa v1 annotation and FaRR1 annotation revealed a substantial improvement in the quality of octoploid gene annotations (Fig 5). We attributed this to the introduction and prioritization of full-length transcript sequences as evidence, in addition to stricter exclusion of gene models encoding smaller proteins, which we were more likely to treat as gene fragments.

**Table 2.**
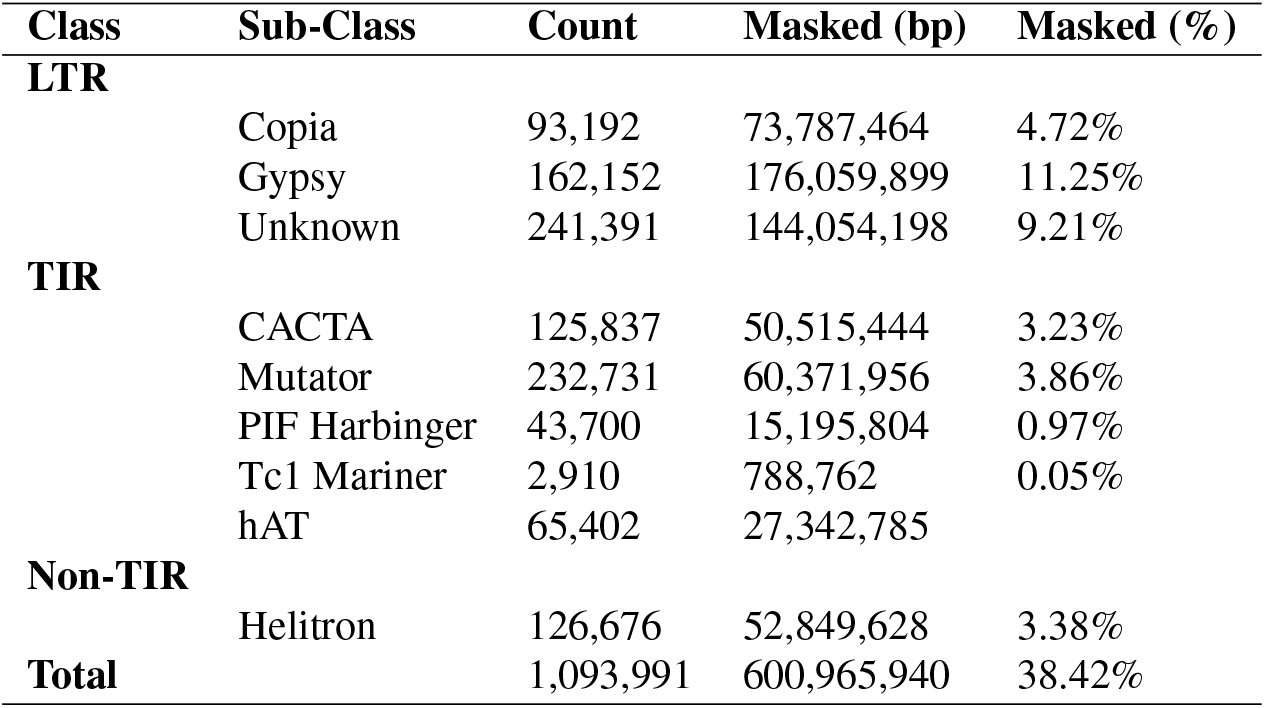
Summary of repetitive DNA elements identified in the FaRR1 genome assembly by “The Extensive *de novo* TE Annotator” pipeline.

| Class | Sub-Class | Count | Masked (bp) | Masked (%) |
| --- | --- | --- | --- | --- |
| LTR | Copia | 93,192 | 73,787,464 | 4.72% |
|  | Gypsy | 162,152 | 176,059,899 | 11.25% |
|  | Unknown | 241,391 | 144,054,198 | 9.21% |
| TIR | CACTA | 125,837 | 50,515,444 | 3.23% |
|  | Mutator | 232,731 | 60,371,956 | 3.86% |
|  | PIF Harbinger | 43,700 | 15,195,804 | 0.97% |
|  | Tc1 Mariner | 2,910 | 788,762 | 0.05% |
|  | hAT | 65,402 | 27,342,785 |  |
| Non-TIR | Helitron | 126,676 | 52,849,628 | 3.38% |
| Total |  | 1,093,991 | 600,965,940 | 38.42% |

**Table 3.** LTR Assembly Index (LAI) scores for ‘Royal Royce’ phase-1 and phase-2 subgenomes, Camarosa v1 octoploid assembly, and FvH4.1 diploid assembly.

| Genome Assembly | Subgenome | Haplotype Phase | LAI Score |
| --- | --- | --- | --- |
| Royal Royce | A | phase-1 | 19.22 |
| Royal Royce | A | phase-2 | 19.08 |
| Royal Royce | B | phase-1 | 18.20 |
| Royal Royce | B | phase-2 | 19.33 |
| Royal Royce | C | phase-1 | 18.83 |
| Royal Royce | C | phase-2 | 18.96 |
| Royal Royce | D | phase-1 | 18.68 |
| Royal Royce | D | phase-2 | 18.76 |
| Camarosa v1 | 1 | na | 12.80 |
| Camarosa v1 | 2 | na | 12.90 |
| Camarosa v1 | 3 | na | 11.66 |
| Camarosa v1 | 4 | na | 13.56 |
| *F. vesca* Hawaii-4 | na | na | 18.44 |

**Fig. 5.**
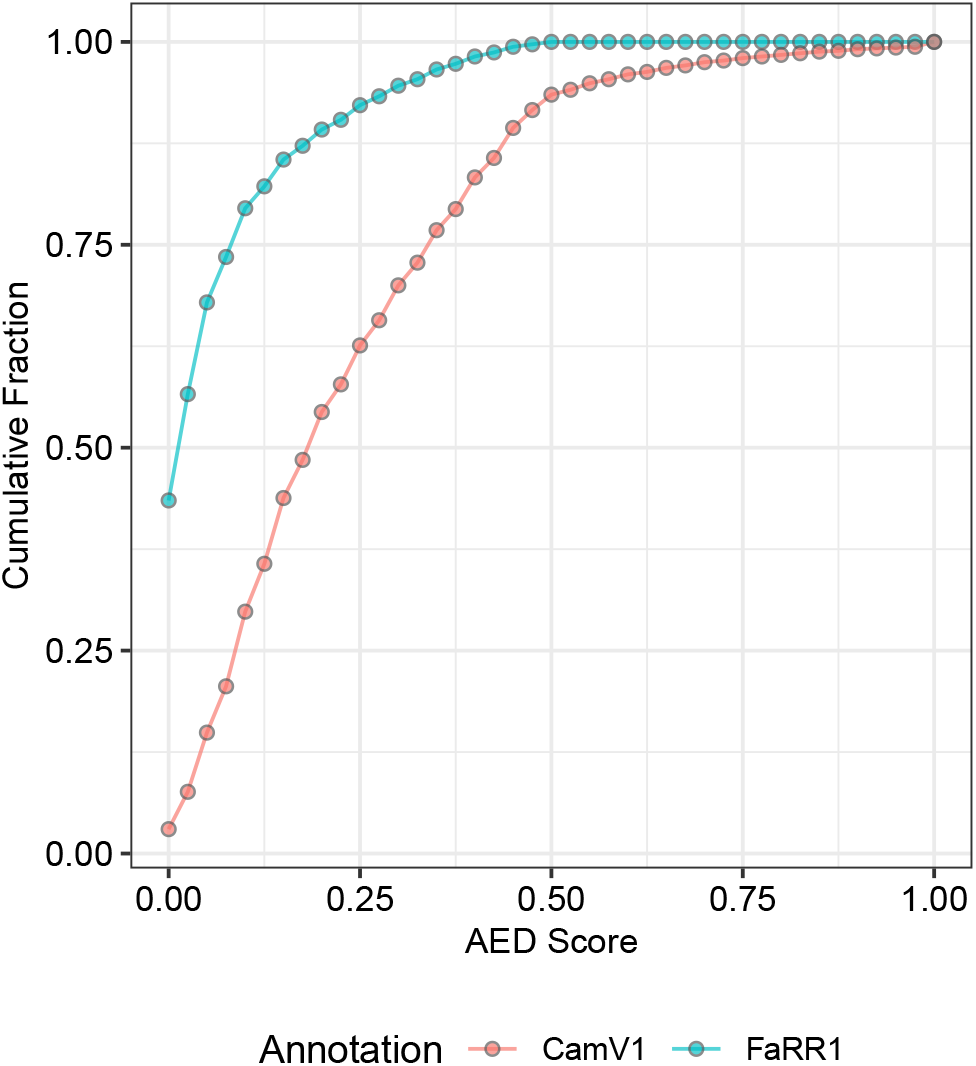
Cumulative annotation edit distance (AED) curves showing the quality of the FaRR1 v1.0 and Camarosa v1 gene annotations based on the cumulative fraction of gene models below a given AED score.

## Conclusions

Our genome assembly and annotation of the heterozygous day-neutral strawberry cultivar ‘Royal Royce’ is a significant improvement on the current publicly available assemblies of octoploid strawberry. The use of HiFi data combining the length of traditional long read sequences with the base-calling accuracy of Illumina short-read sequences helped to minimize the incidence of subgenome chimeras in assembled contigs, and produced highly contiguous assemblies. Due to trio-binning of the HiFi reads using parental sequence information, we also expect that FaRR1 has virtually eliminated the occurrence of phase-switching within the haploid pseudomolecules. By employing dense recombination maps that contain millions of genetically linked variants, saturating physical octoploid haploblocks with parent linkage group information, we were able to manually split the few chimeric assemblies that did occur, confidently bin contigs by chromosome of origin to eliminate spurious allelic contigs, and scaffold the final sets to produce high-confidence pseudo-molecules. Our reassignment of pseudomolecule chromo-some names based on a nomenclature that represents the theoretical diploid origins of the respective subgenomes will facilitate the interpretation of octoploid genetic findings in an evolutionary context, accounting for subgenome dominance and its impact on gene expression and deleterious mutation load. Improvement of gene models will provide strawberry research studies with greater confidence in the structure of genes under investigation. Our re-annotation of the octoploid strawberry genome using IsoSeq data and an expanded pool of transcript evidence has significantly improved the quality of gene models in FaRR1 and the respective phase-1 and phase-2 ‘Royal Royce’ pseudomolecules. With multiple chromosomes represented by a single contiguous sequence, the FaRR1 pseudomolecules are the closest representations of true chromosomes as they exist in an octoploid strawberry nucleus to-date, and offer the most confident representation of the respective octoploid subgenomes.

## Materials and Methods

### Royal Royce HiFi Dataset

We obtained PacBio HiFi sequences for *F. × ananassa* cultivar ‘Royal Royce’ from the dataset published by (6). The HiFi data was generated on the Sequel II platform and contained 83 Gb of sequence in 3.8 M reads. The average read length was 22 kb, with mean RQ value of 28.

### Genomic DNA Sequencing

We performed Illumina whole-genome shotgun sequencing of the University of California, Davis advanced selections ‘04C009P005’ and ‘05C099P006’. Immature leaf tissue was harvested from field grown plants at Wolfskill Experimental Orchard in July of 2019. Leaves were flash frozen, then lyophilized with a Benchtop Pro (VirTis SP Scientific, Stone Bridge, NY) at -40^*°*^C, 100mTorr for 48 hours. For genomic DNA extraction, 30 mg of tissue was ground to a fine powder using steel beads on a Mini 1600 (Spex Sample Prep, Metuchen, NJ). DNA was extracted with a E-Z 96 Plant DNA Kit (Omega Bio-Tek, Norcross, GA, USA) with modifications to increase yield and quality (5). DNA samples were submitted for library preparation and Whole Genome Shotgun Sequencing to the DNA Technologies Core at the Genome Biology Sequencing Facility at the University of California, Davis. DNA samples were submitted to the University of California, Davis DNA Technology Core facility for Illumina 150 bp paired-end library preparation and sequencing on the NovaSeq S4 platform.

### Full Length Transcript Sequencing

We generated PacBio IsoSeq libraries for 15 ‘Royal Royce’ tissues and treatments (Supplementary Table 1). All fruit and nontreated leaf tissue samples were harvested from field grown ‘Royal Royce’ plants in Prunedale, CA between 10:30 AM and 12:00AM on June 9th, 2020. Roughly 20-30 grams of 7 fruit developmental stages were collected from 6 plants: flowers (sepals, petals, calyx), small green fruit, large green fruit, 100% white fruits, partial red fruit, fully red fruit, overripe fruit. Stems were removed from flowers and stems and sepals were removed from fruits and were immediately cut into small pieces and flash frozen in liquid nitrogen. Roughly 10-15 grams of two leaf developmental stages were collected: young, newly emergent leaves and mature leaves. Leaves were flash frozen whole in liquid nitrogen. Samples were transported on dry ice to a -80C freezer before being ground in liquid nitrogen with mortar and pestle. Root tissue was collected from greenhouse-grown ‘Royal Royce’ plants on June 25th, 2020. Two plants placed in a growth chamber were treated with methyl jasmonate on June 23rd, 2020 and roughly 10 grams of young leaf tissue were harvested after 48 h and flash frozen in liquid nitrogen. Roughly 2 grams ground tissue was mixed well with 6 ml warm extraction buffer (2% v/v CTAB, 2% PVP, 100mM Tris, 40% NaCl, 25 mM EDTA, .2g Spermidine, 2% mercaptoethanol) and incubated in a 65C water bath for 5 minutes. Equal volume of 24:1 Chloroform:IsoamylAlcohol was added, mixed well and spun at 4000 rpm for 30 min at 4C; this was repeated using the supernatant. To the supernatant, 10% 1M Potassium Acetate was added, mixed, and spun at 4000 rpm for 20 min at 4C. The resulting supernatant was added to a quarter volume of 10M Lithium Chloride, mixed gently, and precipitated overnight at -20C before being spun at 4000 rpm for 45 min at 4C. The resulting pellet was resuspended in 100 uL RNAse free water and cleaned using the Zymo Research Quick-RNA Mini-Prep Kit and DNase1 treatment (Cat # R1054, Zymo Research Corporation, Irvine, CA) per manufacturer’s instructions. The fifteen ‘Royal Royce’ tissue and treatment RNA samples were submitted to the University of California, Davis DNA Technology Core facility for IsoSeq library preparation. Libraries were pooled and sequenced on two SMRT-cells on the Sequel II platform.

### RNA-seq Datasets

We downloaded sequences from 67 octoploid strawberry RNA-seq libraries representing various *F*. × *ananassa* tissues and treatments from the NCBI sequence read archive (SRA) (https://www.ncbi.nlm.nih.gov/sra) (Supplementary Table 2).

### Trio Binning

We used 150 bp paired-end Illumina WGS sequence datasets from ‘Royal Royce’ maternal parent ‘04C009P005’ and paternal parent ‘05C099P006’ to perform trio-binning of the ‘Royal Royce’ HiFi sequences. We used the scripts published by (7) (https://github.com/esrice/trio_binning) to identify all unique (parent-specific) 21-nt k-mers in the ‘04C009P005’ and ‘05C099P006’ short-read sequences, and produce a histogram of parental k-mer counts. Based on the histogram, we selected a minimum threshold of 20 observations to exclude k-mers representing sequence errors, and a maximum threshold of 250 observations to exclude k-mers representing repetitive sequences. We scanned the ‘Royal Royce’ HiFi sequences for parent-specific k-mers with 20-250 observations and calculated the parental frequencies of unique k-mers in each read. Reads containing at least 100 unique k-mers and at least 80% of k-mers derived from one parent were placed in the corresponding parental read bin. Sequences failing to meet these criteria were assumed to represent fully or mostly homozygous genome sequences and assigned to both parental read bins.

### FALCON Assembly

The two ‘Royal Royce’ HiFi read bins that resulted from trio binning were assembled separately using FALCON (https://github.com/PacificBiosciences/FALCON). We provided FASTQ files as input. We instructed FALCON to ignore error correction by designating `input_type = preads` within the configuration file. We modified the configuration file to provide the following metrics for Daligner to perform pread overlap detection: `ovlp_daligner_option = -e.995 -k24 -h1024 -l5000 -s100`. We modified the configuration file to provide the following metrics for FALCON to perform overlap filtering and assembly: `overlap_filtering_setting = --max_diff 275 --max_cov 275 --min_cov 4 --bestn 15 --n_core 24 --min-idt 99.80 --ignore-indels`. Both assemblies were run on a single node on the FARM computing cluster at the University of California, Davis.

### Recombination Map Anchoring

We obtained the recombination breakpoint maps published by (2), which included over 7,000 total haploblocks containing information from 3.6 M mapped and subgenome-anchored SNP and indel variant sites from Camarosa v1, with corresponding linkage group and genetic map positions. We used BEDTools (22) to extract the 2 kb flanking sequence information from the 3.6 M variant sites in the map, while retaining the linkage group and genetic positions of the variants as identifiers for the resulting markers. We used BLAST to identify the best alignment location of each marker in the phase-1 and phase-2 FALCON contigs, using the center of the alignment as a location to physical anchor the genetically mapped marker. The result was a visual guide of linkage group (chromosome and subgenome) origin and genetic position plotted along each contig’s physical space.

### Subgenome Chimeras and Homologous Allelic Artifacts

We used the information produced by anchoring the recombination breakpoint maps to the phase-1 and phase-2 FALCON contigs as input for the software ALLMAPS (9). We used ALLMAPS to generate warnings for contigs containing mappings from multiple linkage groups. In each of the chimeric contigs flagged by ALLMAPS, we used the input linkage group anchoring information to visually scan the physical contig space for breakpoints between different linkage groups, or non-adjacent map positions within a single linkage group. We used the BEDTools ‘getfasta’ function to “split” the chimeric assemblies at these breakpoints, using the outermost mapped variant sites from the respective independent linkage group regions as borders for the new contigs. We used the dominant linkage group from each contig with physical-genetic marker mappings to bin the respective phase-1 and phase-2 FALCON contig sets into 28 chromosome-specific contig bins. Within each of the 56 chromosome-specific contig bins, we used the function built into the software HaploMerger2 (8) to collapse any contigs fully contained within a larger contig (i.e., redundant) above a minimum identity threshold. We required a minimum identity of 99% for collapsing allelic artifacts.

### Assembly Polishing

The anchoring information for the phase-1 and phase-2 FALCON contigs and the recombination breakpoint maps was used again as input for ALLMAPS to order and orient the contigs, and join using 50 bp sequence gaps. The resulting gapped pseudomolecules were provided as input along with HiFi sequence reads for the software LR_Gapcloser (10), which was running using default parameters to perform gap-filling in the scaffolded assemblies. We used the software Pilon (11) to perform consensus-based error correction of the respective phase-1 and phase-2 assemblies using alignments of their respective trio binned HiFi sequences, and corresponding parental Illumina sequences. Read alignments were performed using BWA-MEM (23) with default parameters. Illumina read alignments with more than a single mismatch were excluded. Pilon was run using default parameters. We aligned the polished pseudo-molecules to the *F. vesca* v4.0 genome assembly (12), and reversed the direction of any pseudomolecules that were not oriented identically to their corresponding diploid chromosome.

### Assembly Quality Assessment

A BUSCO analysis was performed using default parameters to assess the ‘core eudicot’ gene set. The analysis was performed for FaRR1, its four respective subgenomes, and the diploid *F. vesca* Hawaii 4 assembly (12). The LTR Assembly Index (15) for each genome was calculated using whole-genome TE annotations and intact LTR retrotransposons identified by EDTA (24) (https://github.com/oushujun/EDTA), and performed for the four respective subgenomes of ‘Royal Royce’ phase-1 and phase-2 and Camarosa v1, and the complete Hawaii 4 genome (12).

### Transcriptome Assembly

The ‘Royal Royce’ IsoSeq reads were aligned to the assemblies using minimap v2.17 (25) with recommended parameters for spliced long-read alignment: `minimap2 -ax splice:hq –uf <farr1.fa> <isoseq.fq>`. RNA-seq reads were aligned to the assemblies using HISAT v2.2.1 (26) with default parameters. We performed reference-guided transcriptome assembly using StringTie v2.1.4 (16) and the IsoSeq and RNA-seq alignments as input. We provided a GTF guide file (-G) containing spliced alignments of ‘Camarosa v1’ transcripts over 1000 bp in length that aligned to FaRR1 with *≥* 95% coverage and *≥* 98% identity. Otherwise, StringTie was run using default parameters, with the addition of longread (-L) mode for IsoSeq alignments. We used the Mikado v2 (17) pipeline to generate a polished, non-redundant set of phase-1 and phase-2 transcript assemblies that retained the best-scoring transcript evidence at each locus. We used BLASTX to obtain scores for all StringTie assemblies aligned against the UniProt protein database, used TransDecoder v5.5.0 (20) to predict the best six-frame translations of the StringTie assemblies, and used Portcullis v1.2.2 (18) with all merged RNA-seq alignments to predict splice junctions. The Mikado scoring file was modified to prioritize any transcript assemblies derived from IsoSeq alignments over RNA-seq alignments. The resulting files were provided as hints for the Mikado pipeline, in conjunction with the StringTie transcript assemblies. The non-redundant, polished transcripts generated by Mikado were further filtered by re-running TransDecoder to obtain best ORF scores and using HMMER v3.3 (https://github.com/EddyRivasLab/hmmer) to identify associated PFAM domains. Polished transcripts over 200 nt in length, with ORF likelihood scores *≥* 20 and non-TE associated PFAM domains were retained as evidence for gene annotation.

### Custom Repeat Library

We generated the *F. × ananassa* custom repeat library using “The Extensive *de novo* TE Annotator” pipeline (EDTA) (24) (https://github.com/oushujun/EDTA). The pipeline integrates and filters the results of multiple transposable element annotation tools including LTR_FINDER, LTRharvest, Generic Repeat Finder, Helitron Scanner, and TIR-Learner to produce a non-redundant *de novo* TE library. We ran EDTA with default parameters.

### Gene Annotation

We annotated the FaRR1 genome assembly using the MAKER v3.3 annotation pipeline (21). The *F. × ananassa* repeat library generated using EDTA was used to mask the genome assemblies. In the first round of annotation, we provided the 160,375 polished transcript assemblies generated by Mikado as EST evidence. We provided the UniProt plant protein database (19) as protein evidence. The options `est2genome=1` and `protein2genome=1` were used to cause MAKER to output evidence-derived gene models for training SNAP v2013-11-29 (27) and AUGUSTUS v3.2.2 (28). Evidence-derived gene models encoding proteins longer than 75 amino acid residues and with AED scores below 0.20 were used to train SNAP as described by (21), and to train AUGUSTUS as described by (29) before re-running MAKER to predict genes using only the resulting SNAP and AUGUSTUS HMMs. We performed a second round of training and gene prediction with SNAP and AUGUSTUS. The resulting annotations were filtered to remove TE-related predictions and low-quality predictions using the methods described in (29), and corresponding scripts published by their study (https://github.com/Childs-Lab/GC_specific_MAKER). Gene models that encoded proteins with fewer than 65 amino acid residues and had AED scores greater than 0.70 were treated as fragments and excluded.

## Data Access

The FaRR1 genome assembly, gene structural and functional annotations, transcript and peptide sequences, custom repeat library, and supplementary files have been deposited in the DRYAD database (https://doi.org/10.25338/B8TP7G). ‘Royal Royce’ HiFi genomic sequences previously published by (6) are located in the NCBI SRA (Accession: SAMN14691544). ‘Royal Royce’ Iso-Seq transcript sequences and parental Illumina WGS sequences generated for this study are located in the NCBI SRA (BioProject ID: PRJNA765054).

## Bibliography

1. Patrick P Edger, Thomas J Poorten, Robert VanBuren, Michael A Hardigan, Marivi Colle, Michael R McKain, Ronald D Smith, Scott J Teresi, Andrew DL Nelson, Ching Man Wai, et al. Origin and evolution of the octoploid strawberry genome. Nature genetics, 51(3): 541–547, 2019.

2. Michael A Hardigan, Mitchell J Feldmann, Anne Lorant, Kevin A Bird, Randi Famula, Charlotte Acharya, Glenn Cole, Patrick P Edger, and Steven J Knapp. Genome synteny has been conserved among the octoploid progenitors of cultivated strawberry over millions of years of evolution. Frontiers in plant science, 10:1789, 2020.

3. Michael A Hardigan, Anne Lorant, Dominique DA Pincot, Mitchell J Feldmann, Randi A Famula, Charlotte B Acharya, Seonghee Lee, Sujeet Verma, Vance M Whitaker, Nahla Bassil, et al. Unraveling the complex hybrid ancestry and domestication history of cultivated strawberry. Molecular biology and evolution, 38(6):2285–2305, 2021.

4. Dominique DA Pincot, Mirko Ledda, Mitchell J Feldmann, Michael A Hardigan, Thomas J Poorten, Daniel E Runcie, Christopher Heffelfinger, Stephen L Dellaporta, Glenn S Cole, and Steven J Knapp. Social network analysis of the genealogy of strawberry: retracing the wild roots of heirloom and modern cultivars. G3, 11(3):jkab015, 2021.

5. Dominique DA Pincot, Michael A Hardigan, Glenn S Cole, Randi A Famula, Peter M Henry, Thomas R Gordon, and Steven J Knapp. Accuracy of genomic selection and long-term genetic gain for resistance to verticillium wilt in strawberry. The Plant Genome, 13(3):e20054, 2020.

6. Ting Hon, Kristin Mars, Greg Young, Yu-Chih Tsai, Joseph W Karalius, Jane M Landolin, Nicholas Maurer, David Kudrna, Michael A Hardigan, Cynthia C Steiner, et al. Highly accurate long-read hifi sequencing data for five complex genomes. Scientific data, 7(1):1–11, 2020.

7. Sergey Koren, Arang Rhie, Brian P Walenz, Alexander T Dilthey, Derek M Bickhart, Sarah B Kingan, Stefan Hiendleder, John L Williams, Timothy PL Smith, and Adam M Phillippy. De novo assembly of haplotype-resolved genomes with trio binning. Nature biotechnology, 36 (12):1174–1182, 2018.

8. Shengfeng Huang, Mingjing Kang, and Anlong Xu. Haplomerger2: rebuilding both haploid sub-assemblies from high-heterozygosity diploid genome assembly. Bioinformatics, 33(16): 2577–2579, 2017.

9. Haibao Tang, Xingtan Zhang, Chenyong Miao, Jisen Zhang, Ray Ming, James C Schnable, Patrick S Schnable, Eric Lyons, and Jianguo Lu. Allmaps: robust scaffold ordering based on multiple maps. Genome biology, 16(1):1–15, 2015.

10. Gui-Cai Xu, Tian-Jun Xu, Rui Zhu, Yan Zhang, Shang-Qi Li, Hong-Wei Wang, and Jiong-Tang Li. Lr_gapcloser: a tiling path-based gap closer that uses long reads to complete genome assembly. Gigascience, 8(1):giy157, 2019.

11. Bruce J Walker, Thomas Abeel, Terrance Shea, Margaret Priest, Amr Abouelliel, Sharadha Sakthikumar, Christina A Cuomo, Qiandong Zeng, Jennifer Wortman, Sarah K Young, et al. Pilon: an integrated tool for comprehensive microbial variant detection and genome assembly improvement. PloS one, 9(11):e112963, 2014.

12. Patrick P Edger, Robert VanBuren, Marivi Colle, Thomas J Poorten, Ching Man Wai, Chad E Niederhuth, Elizabeth I Alger, Shujun Ou, Charlotte B Acharya, Jie Wang, et al. Single-molecule sequencing and optical mapping yields an improved genome of woodland strawberry (fragaria vesca) with chromosome-scale contiguity. Gigascience, 7(2):gix124, 2018.

13. Jahn Davik, Daniel James Sargent, May Bente Brurberg, Sigbjørn Lien, Matthew Kent, and Muath Alsheikh. A ddrad based linkage map of the cultivated strawberry, fragaria xananassa. PLoS One, 10(9):e0137746, 2015.

14. Felipe A Simão, Robert M Waterhouse, Panagiotis Ioannidis, Evgenia V Kriventseva, and Evgeny M Zdobnov. Busco: assessing genome assembly and annotation completeness with single-copy orthologs. Bioinformatics, 31(19):3210–3212, 2015.

15. Shujun Ou, Jinfeng Chen, and Ning Jiang. Assessing genome assembly quality using the ltr assembly index (lai). Nucleic Acids Research, 46(21):e126, 2018.

16. Sam Kovaka, Aleksey V Zimin, Geo M Pertea, Roham Razaghi, Steven L Salzberg, and Mihaela Pertea. Transcriptome assembly from long-read rna-seq alignments with stringtie2. Genome biology, 20(1):1–13, 2019.

17. Luca Venturini, Shabhonam Caim, Gemy George Kaithakottil, Daniel Lee Mapleson, and David Swarbreck. Leveraging multiple transcriptome assembly methods for improved gene structure annotation. GigaScience, 7(8):giy093, 2018.

18. Daniel Mapleson, Luca Venturini, Gemy Kaithakottil, and David Swarbreck. Efficient and accurate detection of splice junctions from rna-seq with portcullis. GigaScience, 7(12): giy131, 2018.

19. UniProt Consortium. Uniprot: a worldwide hub of protein knowledge. Nucleic acids research, 47(D1):D506–D515, 2019.

20. Brian J Haas, Alexie Papanicolaou, Moran Yassour, Manfred Grabherr, Philip D Blood, Joshua Bowden, Matthew Brian Couger, David Eccles, Bo Li, Matthias Lieber, et al. De novo transcript sequence reconstruction from rna-seq using the trinity platform for reference generation and analysis. Nature protocols, 8(8):1494–1512, 2013.

21. Michael S Campbell, Carson Holt, Barry Moore, and Mark Yandell. Genome annotation and curation using maker and maker-p. Current protocols in bioinformatics, 48(1):4–11, 2014.

22. Aaron R Quinlan and Ira M Hall. Bedtools: a flexible suite of utilities for comparing genomic features. Bioinformatics, 26(6):841–842, 2010.

23. Heng Li. Aligning sequence reads, clone sequences and assembly contigs with bwa-mem. arXiv preprint 1303.3997, 2013.

24. Shujun Ou, Weija Su, Yi Liao, Kapeel Chougule, Jireh RA Agda, Adam J Hellinga, Carlos Santiago Blanco Lugo, Tyler A Elliott, Doreen Ware, Thomas Peterson, et al. Benchmarking transposable element annotation methods for creation of a streamlined, comprehensive pipeline. Genome biology, 20(1):1–18, 2019.

25. Heng Li. Minimap2: pairwise alignment for nucleotide sequences. Bioinformatics, 34(18): 3094–3100, 2018.

26. Daehwan Kim, Joseph M Paggi, Chanhee Park, Christopher Bennett, and Steven L Salzberg. Graph-based genome alignment and genotyping with hisat2 and hisat-genotype. Nature biotechnology, 37(8):907–915, 2019.

27. Ian Korf. Gene finding in novel genomes. BMC bioinformatics, 5(1):1–9, 2004.

28. Mario Stanke, Rasmus Steinkamp, Stephan Waack, and Burkhard Morgenstern. Augustus: a web server for gene finding in eukaryotes. Nucleic acids research, 32(suppl_2):W309–W312, 2004.

29. Megan J Bowman, Jane A Pulman, Tiffany L Liu, and Kevin L Childs. A modified gc-specific maker gene annotation method reveals improved and novel gene predictions of high and low gc content in oryza sativa. BMC bioinformatics, 18(1):1–15, 2017.

